# Characterization, and Anti-microbial Properties of Zinc Oxide Nanoparticles Synthesized Using Citrus sinensis (L.) Osbeck Peel Extract

**DOI:** 10.1101/2020.10.13.337022

**Authors:** Dharma Prasad Khanal, Sadikshya Aryal, Samyam Aryal

## Abstract

**Background:** Citrus sinensis (L.) Osbeck peels are usually discarded as wastes; however, they are rich sources of Vitamin C, fibre, and many nutrients including phenolics and flavonoids which are also good antioxidant agents. This study aimed to examine phytochemical composition, antioxidant capabilities, cytotoxicity of C. sinensis (L.) Osbeck peel extract and and to compare the antibacterial activity with zinc nanoparticles of Citrus sinensis (L.) Osbeck peels with its extract. GC-MS analysis of the compounds present in the peels extract of Citrus sinensis (L.) Osbeck was also done.

**Methods:** C. sinensis (L.) Osbeck fruits were collected from Sindhuli district and were taken to National Herbarium and Plant Laboratory, Godawari, Lalitpur for its identification. Extraction was done by maceration in aqueous solvent. Extract was subjected to Phytochemical screening done by color reactions with different reagents, Antioxidant activities of the peel extracts were examined via the 2,2-diphenylpicrylhydrazyl (DPPH) free radical scavenging activity. Total phenolic content and total flavonoid content of the extracts were measured via the Folin-Ciocalteau method and the aluminium chloride colorimetric method, respectively. Cytotoxic activities of the peel extracts were determined by Brine Shrimp Lethality Bioassay. Comparison of antibacterial activity of extract and zinc oxide nanoparticles prepared via green synthesis using C. sinensis (L.) Osbeck peel extracts as reducing agents. Antibacterial activity was tested by Bore well diffusion method.

**Result:** The extractive value of C. sinensis (L.) Osbeck was found to be 8.64% in aqueous solvent. GC-MS analysis of peel extract of C. sinensis (L.) Osbeck showed the presence of 2-Methoxy-4-vinylphenol, 4H-Pyran-4-one, 2,3-dihydro-3,5-dihydroxy-6-methyl, Benzoic acid, 3-Deoxy-d-mannoic lactone and 5-Hydroxymethyl-furfuralas major compounds. The qualitative phytochemical test showed the presence of tannin, alkaloid, carbohydrate, flavonoid, cardiac glycoside, terpinoid.

The DPPH radical scavenging activity of C. sinensis (L.) Osbeck peel extract was 35.56 μg/ml. TPC of C. sinensis (L.) Osbeck peel extracts was 46.07 mg GAE/g. TFC was 1.29 mg QE/g. The LD50 value of Brine Shrimp Lethality assay of the extract showed 312.5μg/ml which is indicative. The antibacterial activity of zinc oxide nanoparticles was found to be greater than that of the extract, but the antibacterial activity of Zn-NPs was less than that of the standard.

**Conclusion:** Hence, the GC-MS analysis of aqueous extracts of leaves of C. sinensis (L.) Osbeck showed the presence of 20 different compounds. Phytochemicals including phenolics and flavonoids in C. sinensis (L.) Osbeck peel extracts exhibited good antioxidant properties. The extract also exhibited antibacterial activity which was 4 times less than that of the standard. The antibacterial activity of standard was 2 times greater than that of Zn-NPs. The extract also exhibited cytotoxic activity. This study indicated that C. sinensis (L.) Osbeck peels contained potential antioxidant, cytotoxic and antibacterial compounds which could be exploited as value added products.

## Introduction

Nanoparticles are materials in a nanoscale ^1^. The diameter of these particles ranges between 1nm and 100 nm. Nanoparticles have found their usage in medicine, agriculture, and food industry. The currently prevalent physiochemical method of preparation of Nanoparticles is hazardous, environmentally unfriendly, expensive, requires condition of high temperature, pH, and/or pressure for synthesis ^2^. As such potential adverse effect on human as well as environmental have often been raised. Biological methods of synthesis of nanoparticles can provide a great ecofriendly and safe alternative provided that it is sustainable and scalable.

Nanoparticles exhibit new and enhanced biochemical properties as well as distinctly improved phenomenon and functionality. Given these size-controlled particles display significantly varying properties as compared to macro-particles, these particles should be studied and characterized. Also, these modified physical, chemical, and morphological properties of Nanoparticles allow for a unique interaction with cellular molecules while facilitating the entry into the inner-cellular compartments ^3^. Also, there is greater surface area and consequently greater reactivity of Nanoparticles when compared to the macro-sized particles.

Green synthesis is economically and ecologically viable way for synthesis of Nanoparticles for various industrial application. It is known that extracts may function as reducing as well as stabilizing agent during the synthesis of NPs. Nobel metals like copper, iron, zinc, silver, gold, silver, platinum, and palladium have been frequently used for the creation of nanoparticles ^4,5^.

The therapeutic and diagnostic application of nanoparticles have received substantial consideration in the recent years. The interaction of nanoparticles and nanoparticles at subcellular levels greatly enhances the signaling of markers in case of diagnostics and improved specificity of targeted therapeutics. Biosynthesized nanoparticles are also the ideal nominees for therapeutic application as they are biologically compatible and naturally safe.

Zinc oxide is a photo catalyst for the degradation of organic pollutants. As such, zinc oxide can be considered one of the candidates for green synthesis of nanoparticles. Combined with this fact that aqueous extracts of *Citrus sinensis* (sweet orange) contains high amount of phenolic compound which could possibly help with reduction process and better synthesis ^6^. Zinc nanoparticles are much cheaper than other nanoparticles – say gold or silver making in much more feasible during synthesis.

Several, study have shown that nanoparticles – especially silver nanoparticles – have a potential antibacterial property. Further studies on silver NPs show that biogenic nanomaterials are biologically compatible and could possibly serve as treatment against infectious diseases ^7^. This anti-microbial property could potentially be evaluated in zinc oxide nanoparticles as well.

Synthesized antibiotics are becoming redundant against several micro-organisms. Plant extracts haven’t found a use in medicine because of several obstacles. Several proposals seek to combine the plant extracts with synthesis of NPs to increase the effectiveness of such extracts.

Phytochemical composition and in vitro antioxidant activities and found DPPH (1,1-Diphenyl-2-Picrylhydrazyl) free radical scavenging activity of *C. sinensis* (*L*.) *Osbeck peel* extracts in various solvents was between 8.35 - 18.20 mg TE/g. TPC of peel extracts, for the same, ranged from 12.08 - 38.24 mg GAE/g. The TFC ranged from 1.90 mg CE/g to 5.51 mg CE/g ^8^.

Also, phytochemical analysis of *Citrus sinensis* (*L*.) *Osbeck* peel and found that the phytochemical analysis of methanolic extract of *C. sinensis* (*L*.) *Osbeck* has shown the existence of secondary metabolites such as tannin, flavonoids, reducing sugar, alkaloids, cardiac glycoside as well as the aqueous extract which showed the same thing ^9^.

Green synthesis of Zinc Oxide nanoparticles using *Citrus sinensis* (*L*.) *Osbeck* extract presents the Zn O bond at 618cm^−1^, a crystalline growth in an exclusively a hexagonal wurtzite crystal structure, and different size and shape homogeneity that was contingent on the amount of extract used ^10^.

Previously conducted analysis on ZnO nanoparticles structure synthesized using *Trifolium pretense* L. flower extract was characterized by UV–Vis spectroscopy, Fourier transform infrared spectroscopy (FT-IR), X-ray diffraction (XRD), and scanning electron microscopy (SEM) with Energy Dispersive X-ray analysis (EDX) ^11^.

Owing to previous such studies, the present studies tries to characterize zinc oxide nanoparticles synthesized with the use of *Citrus sinesis* and also looks into the antimicrobial action of biosynthesized zinc oxide nanoparticles.

## Methodology

### Design

The present study is a combination of descriptive and experimental designs. The study has been approved and conducted under the guidelines of IRC, ManMohan Institute of Health Sciences.

Collection and Identification of Samples -The fruits of *Citrus sinensis* (L.) were collected from Sindhuli, Nepal on January 2019. The preserved herbarium of the plant was submitted to the National Herbarium and Plant Laboratory, Godawari, Nepal for identification.

### Processing the samples

The peels of Citrus sinensis (L.) were washed properly to remove contaminations like dust and soil. The cleaned peels were shade dried for several days and then cut into pieces. The dried peels were then grounded into coarse powder using electric blender. A sieve of 500 microns was used to obtain the fine powder which was then subjected to extraction.

### Extraction

50gm of dried peel powder with 500 ml of distilled water was used for maceration. It was allowed to stand for 3 days with occasional stirring. It was then filtered using muslin cloth and again by Whatman Filter Paper Grade No. 1. The filtrate was evaporated in rotary evaporator and obtained extract was transferred to a stainless steel plate. Then, the extract was concentrated at a room temperature. The dried extract was scraped off the plate and stored in a borosilicate glass vials. The percentage yield of the dried plant extracts was calculated using:

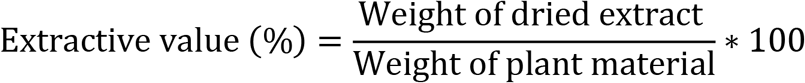

### Preliminary phytochemical screening ^12,13^

Phytochemical screening was carried out to identify the major groups of phytochemicals present in the dried extracts of *Citrus sinensis* (*L*.) Osbeck by various reagents. The extracts were subjected for alkaloids, glycosides, tannins, carbohydrates, total phenolic content^14^, total flavonoid content^15^, saponin, and terpinoid testing with appropriate modifications.

The extract was evaluated using–mass spectrometry (GC–MS)(GCMS-QP2010 Ultra) analysis to identify the chemical compunds present in the extract that showed various biological activities. The column dimension was 30m x 0.25mmID df RTX 5MS Column. Helium was the preferred carrier gas and the temperature programming was set with initial oven temperature at 100°C and held for 1 min and the final temperature of the oven was 300°C with rate at 30C and hold out time of 8 min. A 2μL sample was injected in splitless mode. The mass spectra were recorded over 40.00 m/z to 600.00 m/z. The running time was 20 minutes. The constituents were determined by relating the respective peaks to Total Ion Chromatogram (TIC) areas obtained from the GC-MS.

### Evaluation of antioxidant activity ^16,17^

The radical scavenging activity of extract was based on the comparison with scavenging activity of the stable 1, 1-diphenyl-2-picrylhydrazyl (DPPH) free radical was determined.

Reference samples of ascorbic acid and sample solutions of plant extract were prepared using methanol as base at different concentrations (5, 10, 15, 20 and 25 μg/ml). 0.1 mM solution of DPPH with methanolic base was prepared and 3 ml of this solution was added to 1 ml of each different concentration of sample plant extracts and ascorbic acid solutions. This blend was kept in dark for 30 minutes. Similarly, as control, 3 ml of 0.1 mM DPPH was mixed with 1 ml of methanol (solvent) and kept in dark for 30 minutes. Thirty minutes later, the absorbance was measured at 517nm. The capability to scavenge the DPPH radical was calculated by using the following formula:

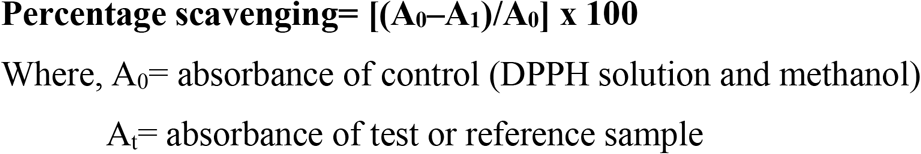

The % scavenging was then plotted against concentration and regression equation was obtained. IC_50_ (microgram concentration required to inhibit DPPH radical formation by 50%) values were calculated for each plant extracts by using this inhibition curve.

Lower absorbance of the reaction mixture indicates higher free radical scavenging activity.

### Preparation of 0.1 mM DPPH in methanol

1 mole is equivalent to 394.32 gm of DPPH

1/1000mM is equivalent to 394.32/1000 mg of DPPH

0.1mm is equivalent to 39.432* 0.1 mg of DPPH

39.432mg of DPPH in 1000ml of methanol

### Brine Shrimp Lethality Bioassay ^9^

The artificial sea water was prepared by dissolving 25gm of NaCl in 1000 ml of water. The cytotoxic activity of the plant was evaluated using Brine shrimp lethality bioassay method where 6 graded doses (*viz*, 800 μg/mL, 400 μg/mL, 200 μg/mL, 100 μg/mL, and 50 μg/mL) were used. Mature brine shrimps (*Artemia salina* Leach) nauplii were used as test organisms. For hatching, eggs were kept in artificial sea salt with a constant oxygen supply for 48 h. Water was used as a solvent and also as a negative control. Vincristine sulfate was used as a reference standard. The numbers of survivors were counted after 24 h. Larvae were considered dead if they did not exhibit any internal or external movement during several seconds of observation. The larvae did not receive food. To ensure that the mortality observed in the bioassay could be attributed to bioactive compounds and not to starvation; we compared the dead larvae in each treatment to the dead larvae in the control.

The median lethal concentration (LC_50_) of the test samples were calculated using the probit analysis method described by Finney^18^, as the measure of toxicity of the plant extract.

### Mortality %= No. of Dead larvae/Total no. of Larvae *100%

#### Synthesis of Zno-NPs ^19,20^

##### Principle

The growth process of ZnO nanoparticles can be controlled through the following listed chemical reactions:

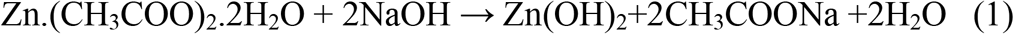

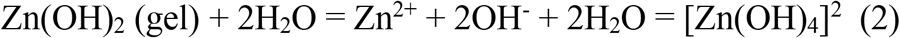

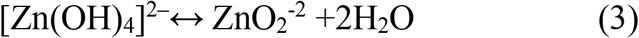

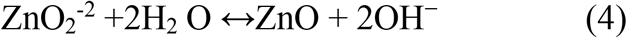

The formation mechanism of the ZnO nanostructures is a complex process and in summary considered to include two main steps: the generation of a ZnO nuclei, and subsequent ZnO crystal growth. The Zn(OH)_4_^2-^ complexes serve as basic growth units for the preparation of ZnO nanostructures. The zinc acetate may convert into Zn(OH)_2_ colloids firstly under alkali solution, as shown in reaction 1. During the process, part of the Zn(OH)_2_ colloids dissolves into Zn^2+^ and OH^−^ according to reaction no 2. When the concentration of Zn^2+^ and OH^−^ reaches the supersaturation degree of ZnO, ZnO nuclei will form according to reaction no 4.

#### Preparation of peels extract of *C. sinensis* (*L*.) *Osbeck* peels

The dried peels of orange (10 g) were boiled in 100ml distilled water for 15 min. The resulting extract was cooled and filtered using Whatman Filter Paper Grade No. 1 and used as the extract solutions.

#### Synthesis of nanoparticles

In this method, 0.02 M solution of zinc acetate (50ml) was taken and 2ml peel extract was added drop-wise and the resulting mixture was stirred for 10 minutes. The pH of the mixture was maintained at 12 by adding 0.02M NaOH drop-wise and the solution was stirred continuously for 2 hr. A pale white precipitate resulted which was washed by distilled water 3 times followed by ethanol and dried at 60°C overnight in oven. Pale white powder of zinc oxide nanoparticles was store for characterization.

### Characterization of Zno-NPs

#### UV-Visible spectroscopy

The formation of zinc -NPs was characterized by measuring the surface plasmon resonance (SPR) of the solution in the range from 200 to 800 nm in UV–Vis spectroscopy ^21^, which has proven to be a useful spectroscopic method for the detection of prepared metallic nanoparticles. The distilled water was used as blank.

#### X-ray diffraction

X-ray diffractions (XRD) was used to affirm the crystal phases and size. It was carried out using X-ray diffractometer of crystal radiation (λ = 1.541 A°) for (2theta) range of (20°-90°). The full widths at half maximum (FWHM) in the XRD was used to determine the crystallite size by Scherer’s equation ^22^:

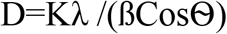

Where, D is the mean size of crystal

K is the crystalline shape factor i.e 0.89
ß is the structural broadnening (FWHM)
θ is the Bragg’s angle.

#### Fourier Transform Infrared Spectroscopy (FT-IR)

FTIR of extract and synthesized nanoparticles was done to identify the functional group present and to single out possible biomolecules responsible for reduction and stabilization of zinc acetate to zinc oxide.

### Anti bacterial activity ^8^

#### Evaluation of anti bacterial activity

Antimicrobial activity was evaluated by agar well method. In the method, test organisms were collected and were standardized with reference to 0.5M MacFarland standard.

#### Microorganism culture

Microorganism susceptible to commercial grade of antibiotics was deemed selectable for the analysis of antimicrobial properties. ATCC culture was collected from Natural Product Research Laboratory, Thapathali, Kathmandu. The organisms under evaluation were:

Gram positive: *Staphylococcus aureus* (ATCC no. 6538P)*, Bacillus subtilis* (ATCC no. 6051)
Gram negative: *Escherichia coli* (ATCC no. 8739)*, Klebsella pneumonia* (ATCC no. 700603)
The collected microorganisms were cultured in freshly prepared petri plates of Mueller Hilton Agar (MHA). Before the test was performed the organism were let to grow in Peptone water broth for the standardization.

#### Standard preparation

The standard used for the antimicrobial evaluation was 320 μg/ml, 160 μg/ml, 80 μg/ml aSnd 40 μg/ml of Azithromycin and Gentamycin solution which was dissolved in DMSO (1%) solution for gram positive and gram negative organisms, respectively.

#### Test procedure ^23^

For the test, agar well diffusion method was used. MHA agar was prepared as per the manufacturer’s instructions. The agar was prepared and sterilized in autoclave at 15lbs pressure for 15 minutes. The agar was poured in sterile petri plates in sterile laminar hood. It was allowed to set. 8mm well borer was used to prepare bores for the test procedure. The microorganisms were swabbed in the petri plates using sterilized swab stick. 100 μl of the extracts were taken with the help of micropipette having sterile tips. 4 holes were bored per plate for 4 different concentrations of the test extracts. Similar procedure was repeated for standard and blank. The plates were left to incubate and the zone of inhibition was determined after 24 hours.

## RESULTS and Discussion

### Extractive Value

4.32gm of extract was obtained from 50 gm of dried peel powder by successive maceration. The extractive value of *Citrus sinensis (L.) Osbeck* peels in distilled water was found to be 8.64%. The extract was brownish in color and sticky mass in nature.
In the previous study by Musa et.al. in 2019 the extractive value was 19% ^8^. The difference in extractive value may be due to geographical variation and the extraction in the article was done by continuous stirring.

### Preliminary Phytochemical Screening

The different phytochemicals in the extract were identified by the chemical reaction and color change with different reagents. Phytochemical analysis aqueous extract of plant was done for presence or absence of secondary metabolites or different constituents such as alkaloids, glycosides, flavonoids, phenol, tannin, carbohydrates, proteins, saponins and amino acids (table 1). Anthraquinones and saponin were absent. Tannins, carbohydrates, flavonoids, alkaloids, terpenoids and cardiac glycosides were present which is similar to the findings research done by Musa et. al ^8^.

**Table 1:** Phytochemical screening of the extract of *C. sinensis* (L.) Osbeck

| Test | Result |
| --- | --- |
| Tannin (Ferric chloride test) | + |
| Alkaloid<br>- Hager's<br>- Dragendorff's<br>- Mayer's<br>- Wagner's | -<br>-<br>+<br>+ |
| Saponin (Frothing test) | - |
| Glycoside (Fehling's test) | + |
| Cardiac glycoside (Keller Killiani test) | + |
| Anthraquinone glycoside<br>(Borntranger's test) | - |
| Carbohydrate (Molisch's test) | + |
| Terpinoid (Salkowski's test) | + |
| Flavonoid<br>- Shinoda test<br>- Alkaline reagent test<br>- Zinc hydrochloride test | +<br>+<br>+ |

Note: (+) indicates presence (-) indicates absence of phytochemicals

### Quantitative Phytochemical Screening

#### Total Phenolic content determination

The total phenolic contents of the extracts as determined by Folin–Ciocalteu method are reported as gallic acid equivalents. The concentration of phenolic in extracts were calculated from the calibration curve using the regression equation Y= 0.0115x-0.0349, R^2^=0.9984 (table 2, figure1). The absorbance values obtained at different concentrations of gallic acid was used for the construction of calibration curve. Absorbance values for gallic acid is measured at 760 nm using Folin-Ciocalteu reagent.

**Table 2:** Absorbance value of gallic acid measured for calibration curve

| Concentration of gallic acid (mcg/ml) | Absorbance for Gallic Acid Measured |
| --- | --- |
| 10 | 0.031±0.023 |
| 25 | 0.277±0.012 |
| 50 | 0.583±0.014 |
| 100 | 1.05±0.014 |
| 200 | 2.265±0.019 |

**Figure 1:**
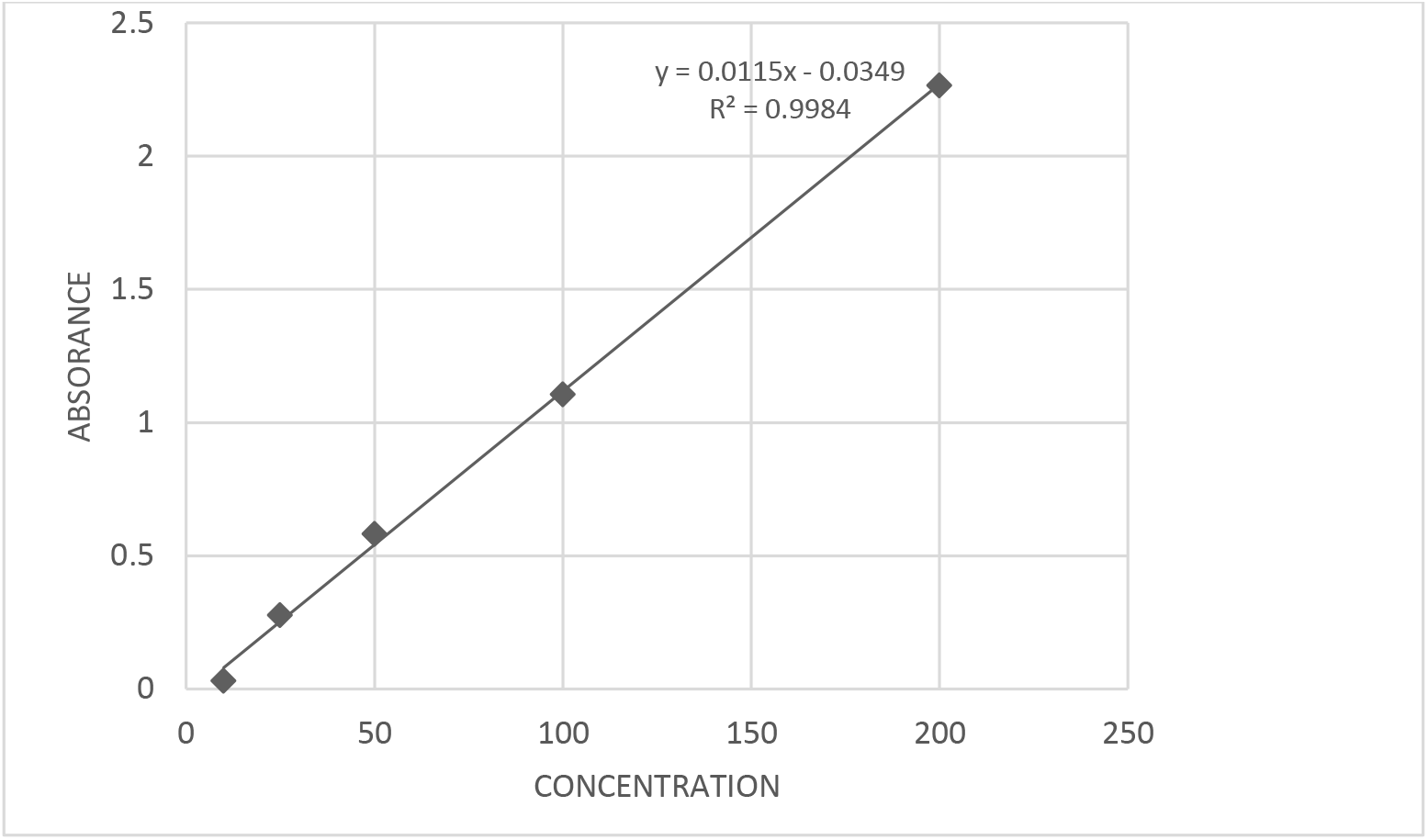
Gallic acid standard calibration curve. The TPC in aqueous extract was found to be 46.07mg GAE/g (table 2). In the previous study by Liew et. al. in 2018, the TPC was found to be 12.08 mg GAE/g ^24^. The difference in the TPC value may be due to different experimental conditions and geographical variation.

**Table 2:** Absorption value of the extract

| Extract | Average Absorbance |
| --- | --- |
| Aqueous | 0.495±0.003 |

#### Total flavonoid content determination

The total flavonoids content of the extracts and essential oil as determined by a colorimetric assay using aluminum chloride and are reported as quercetin equivalents. The concentration of flavonoid in extract was calculated from the calibration curve using the regression equation Y=0.0017x + 0.0248, R^2^ = 0.9710 (table 3, figure 2). The absorbance values obtained at different concentrations of quercetin was used for the construction of calibration curve. Absorbance value for quercetin was measured at 510 nm in UV and is mentioned at Table 4.

**Table 3:** Absorbance value for quercetin measured for calibration curve

| Concentration of quercetin (mcg/ml) | Absorbance for quercetin measured |
| --- | --- |
| 10 | 0.024±0.001 |
| 25 | 0.063±0.004 |
| 50 | 0.146±0.001 |
| 100 | 0.175±0.003 |
| 200 | 0.361±0.0005 |

**Figure 2:**
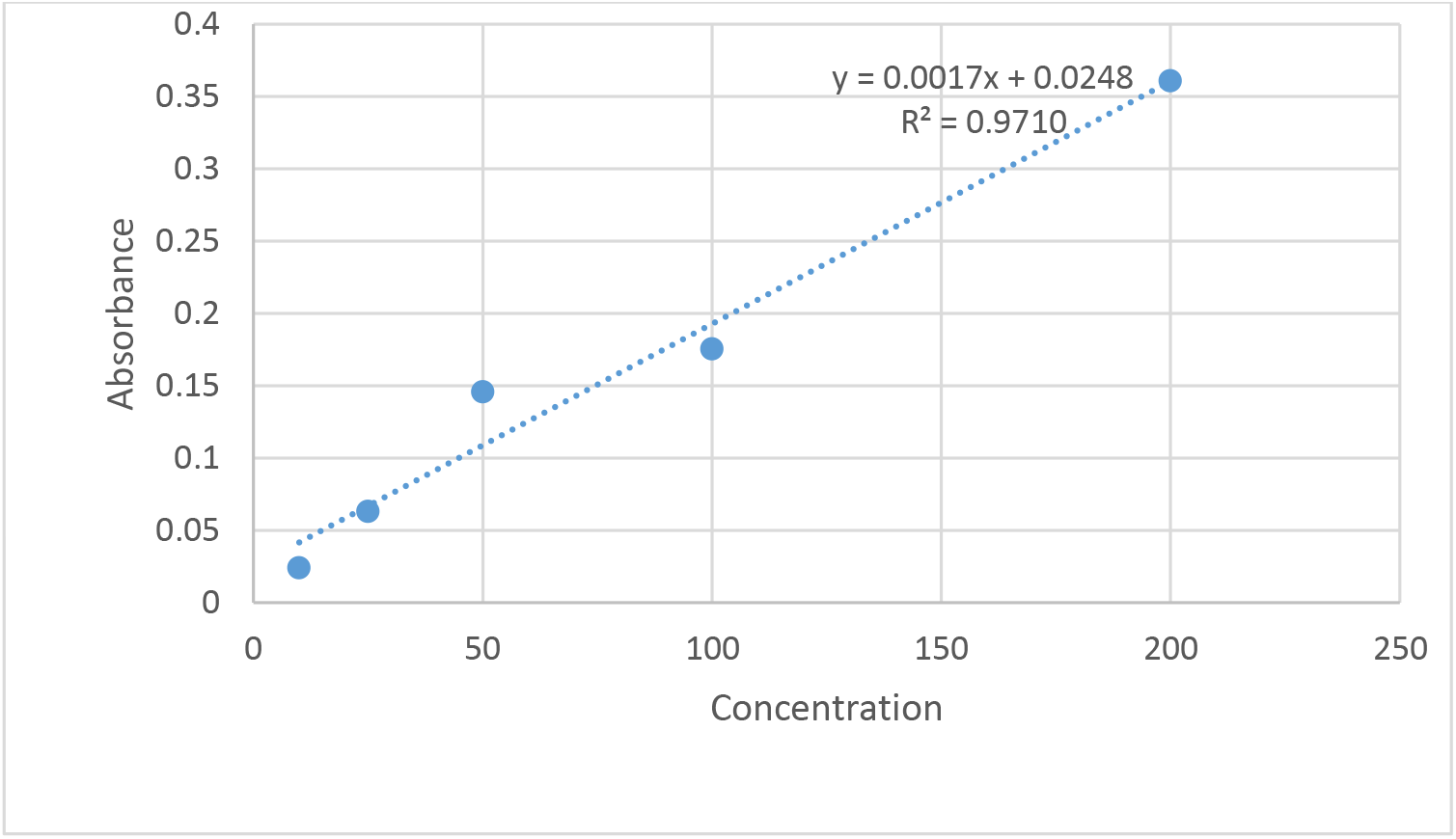
Quercetin standard calibration curve. The TFC value was found to be 1.29 mg QE/g (table 4). The TFC value was found to be 1.26 mg QE/g. In the previous study by Liew et. al. in 2018, the TPC was found to be 1.90 mg QE/g ^24^.

**Table 4:** Absorption Value of extract

| Extract | Absorbance |
| --- | --- |
| Aqueous | 0.027±0.001 |

### GC-MS analysis

GC-MS analysis of the aqueous extract was performed.

**Figure 3:**
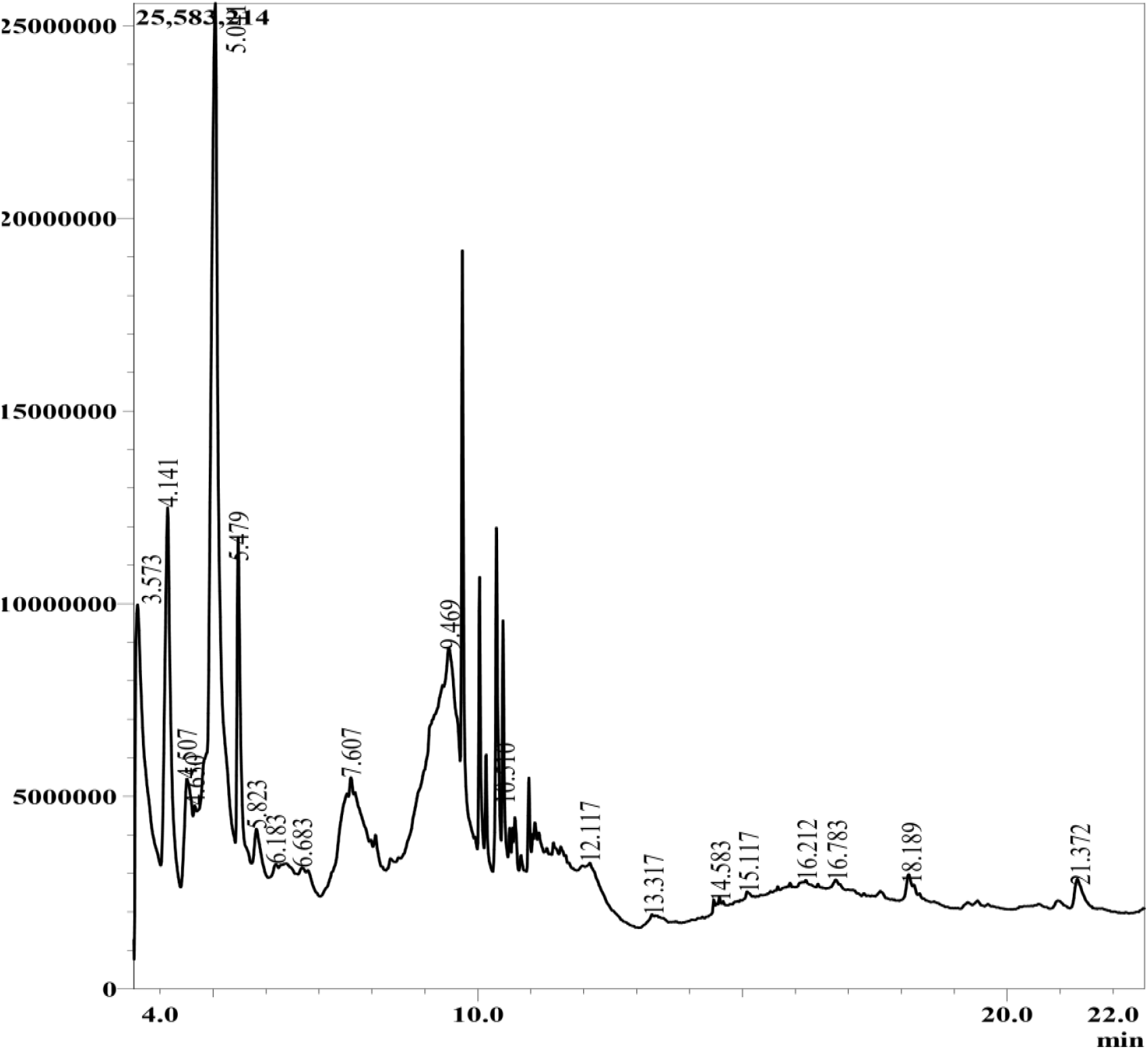
GC-MS Chromatogram of aqueous extract of *C. sinensis* (L.) Osbeck

**Table 5:** Composition of aqueous Extract

| Peak | R. Time | Area % | Name | Base m/z |
| --- | --- | --- | --- | --- |
| 1 | 4.507 | 2.36 | Benzoic acid | 105.05 |
| 2 | 4.650 | 1.32 | 4-Methyleneisophorone | 107.10 |
| 3 | 5.041 | 16.32 | 5-Hydroxymethylfurfural | 41.05 |
| 4 | 5.479 | 3.86 | 2-Methoxy-4-vinylphenol | 150.15 |
| 5 | 5.823 | 2.19 | 4,5-Dihydro-2-methylimidazole-4-one | 42.05 |
| 6 | 6.183 | 3.34 | Ethyl trans-3-methyl-2-oxiranecarboxylate | 45.05 |
| 7 | 6.683 | 2.01 | l-Gala-l-ido-octose | 43.05 |
| 8 | 7.607 | 9.72 | Adenosine, N6-phenylacetic acid | 57.05 |
| 9 | 9.469 | 24.39 | 3-Deoxy-d-mannoic lactone | 44.05 |
| 10 | 10.510 | 11.47 | Dibutyl phthalate | 149.10 |
| 11 | 12.117 | 2.51 | Scyllo-Inositol | 73.05 |
| 12 | 16.212 | 2.84 | 7-Hydroxy-3-(1,1-dimethylprop-2-enyl)coumarin | 57.10 |
| 13 | 21.372 | 1.41 | 3',4',5,6,7,8-Hexamethoxyflavone | 387.10 |

**Table 6:**
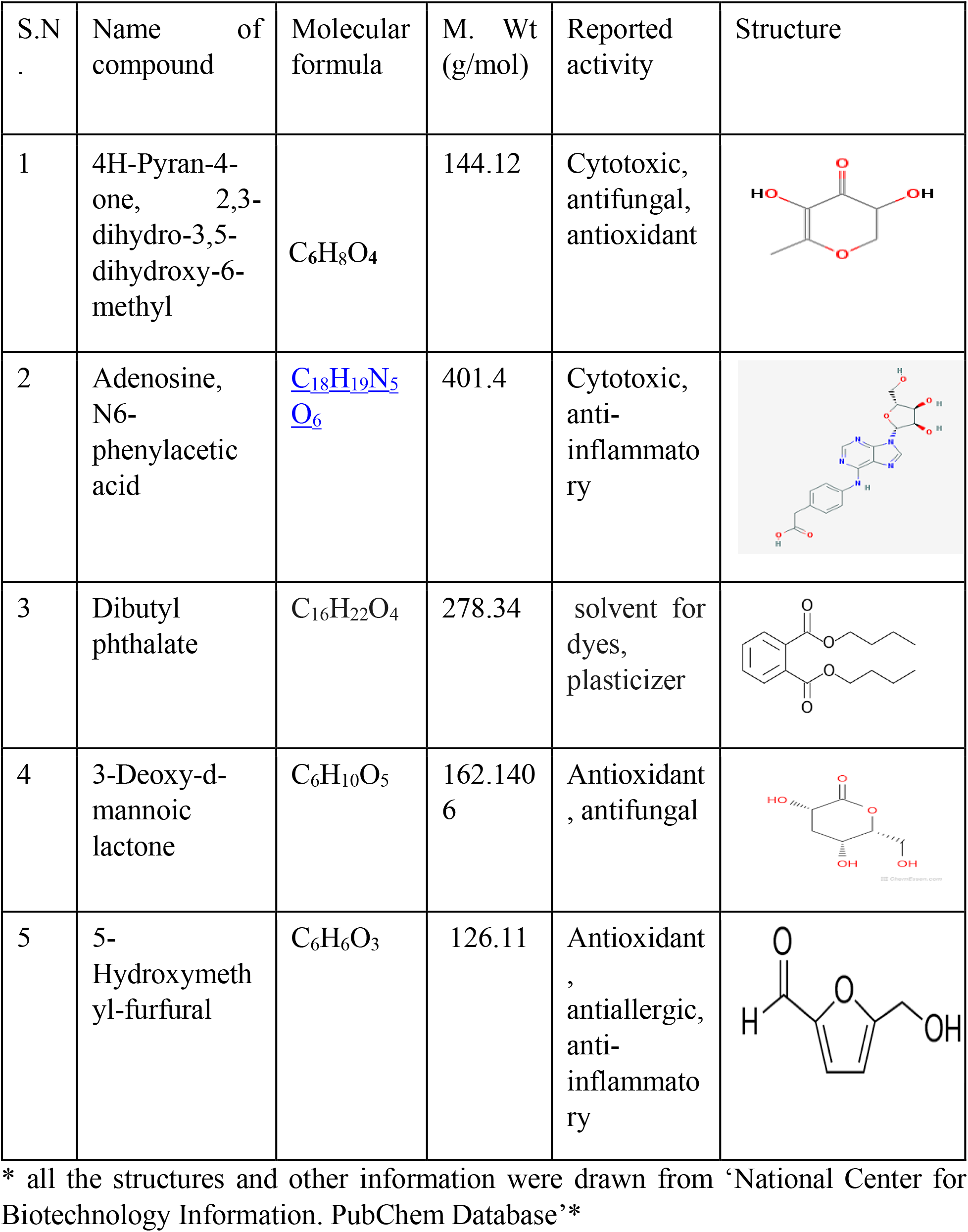
The five major compounds found in the extract

| S.N . | Name of compound | Molecular formula | M. Wt (g/mol) | Reported activity | Structure |
| --- | --- | --- | --- | --- | --- |
| 1 | 4H-Pyran-4-one, 2,3-dihydro-3,5-dihydroxy-6-methyl | $C_6H_8O_4$ | 144.12 | Cytotoxic, antifungal, antioxidant | |
| 2 | Adenosine, N6-phenylacetic acid | $C_{18}H_{19}N_5O_6$ | 401.4 | Cytotoxic, anti-inflammatory |  |
| 3 | Dibutyl phthalate | $C_{16}H_{22}O_4$ | 278.34 | solvent for dyes, plasticizer | |
| 4 | 3-Deoxy-d-mannoic lactone | $C_6H_{10}O_5$ | 162.1406 | Antioxidant, antifungal | |
| 5 | 5-Hydroxymethyl-furfural | $C_6H_6O_3$ | 126.11 | Antioxidant, antiallergic, anti-inflammatory | |

\* all the structures and other information were drawn from ‘National Center for Biotechnology Information. PubChem Database’\*

According to the research done by Oluwole et.al the Gas Chromatography-Mass Spectrometry analysis revealed that ethanolic peel extract of *C. sinensis (L.) Osbeck* contains 5, 6, 7, 8, 3’, 4’-hexamethoxyflavone which is commonly called nobiletin. The aqueous extract also showed the same result ^25^.

### Antioxidant Activity

The antioxidant potential was assessed in terms of IC_50_ by DPPH free radical scavenging capacity. IC_50_ value of extract and standard was calculated through regression equation. The lower IC_50_ value represents higher scavenging abilities. The IC_50_ value of the extract was found to be 35.56 μg/ml. Ascorbic Acid was used as standard and its IC_50_ value was found to be 9.72 μg/ml (table 7).

**Table 7:**
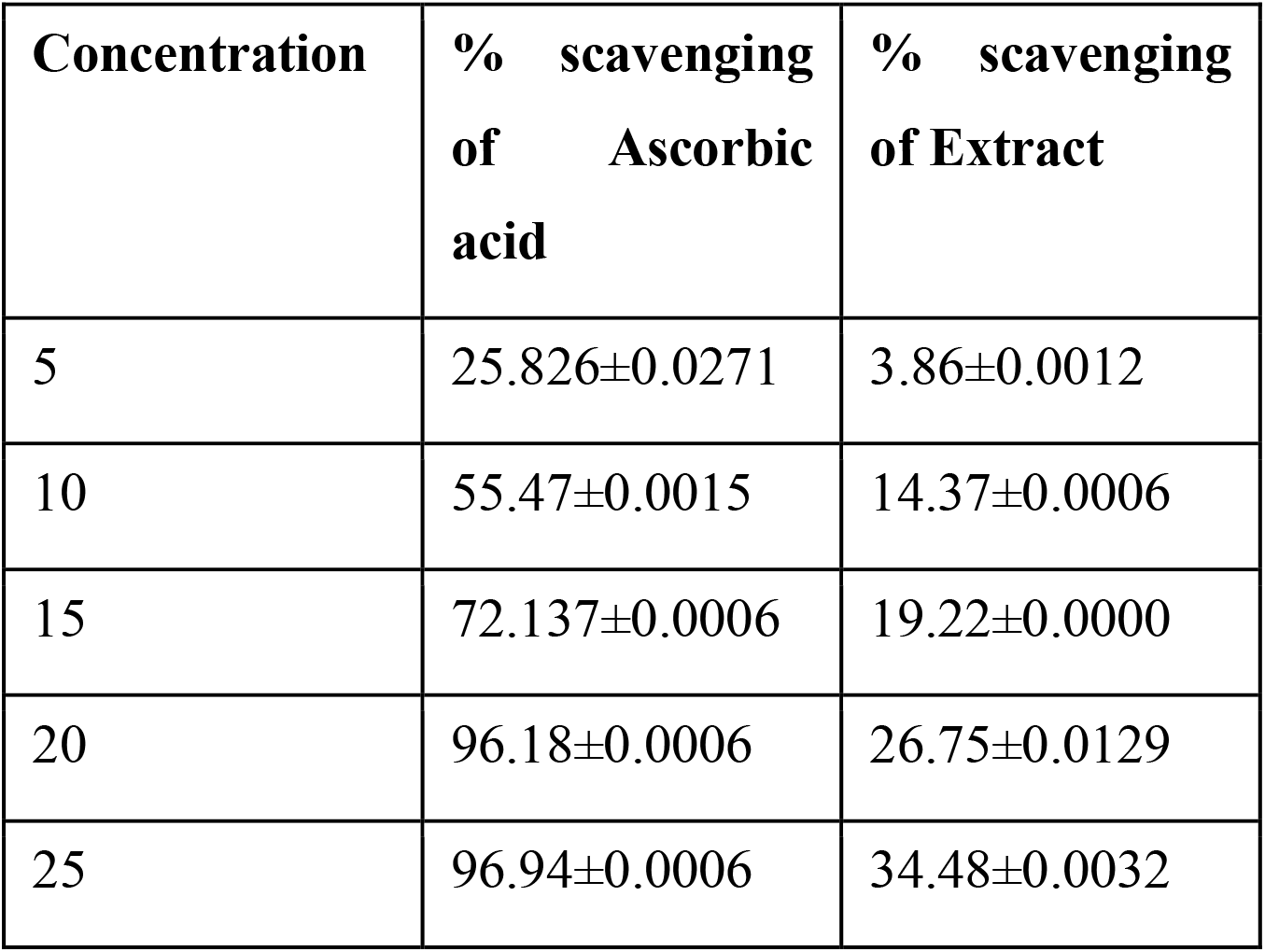
Percentage scavenging of extracts and ascorbic acid at different concentration

| Concentration | % scavenging of Ascorbic acid | % scavenging of Extract |
| --- | --- | --- |
| 5 | 25.826±0.0271 | 3.86±0.0012 |
| 10 | 55.47±0.0015 | 14.37±0.0006 |
| 15 | 72.137±0.0006 | 19.22±0.0000 |
| 20 | 96.18±0.0006 | 26.75±0.0129 |
| 25 | 96.94±0.0006 | 34.48±0.0032 |

**Figure 4:**
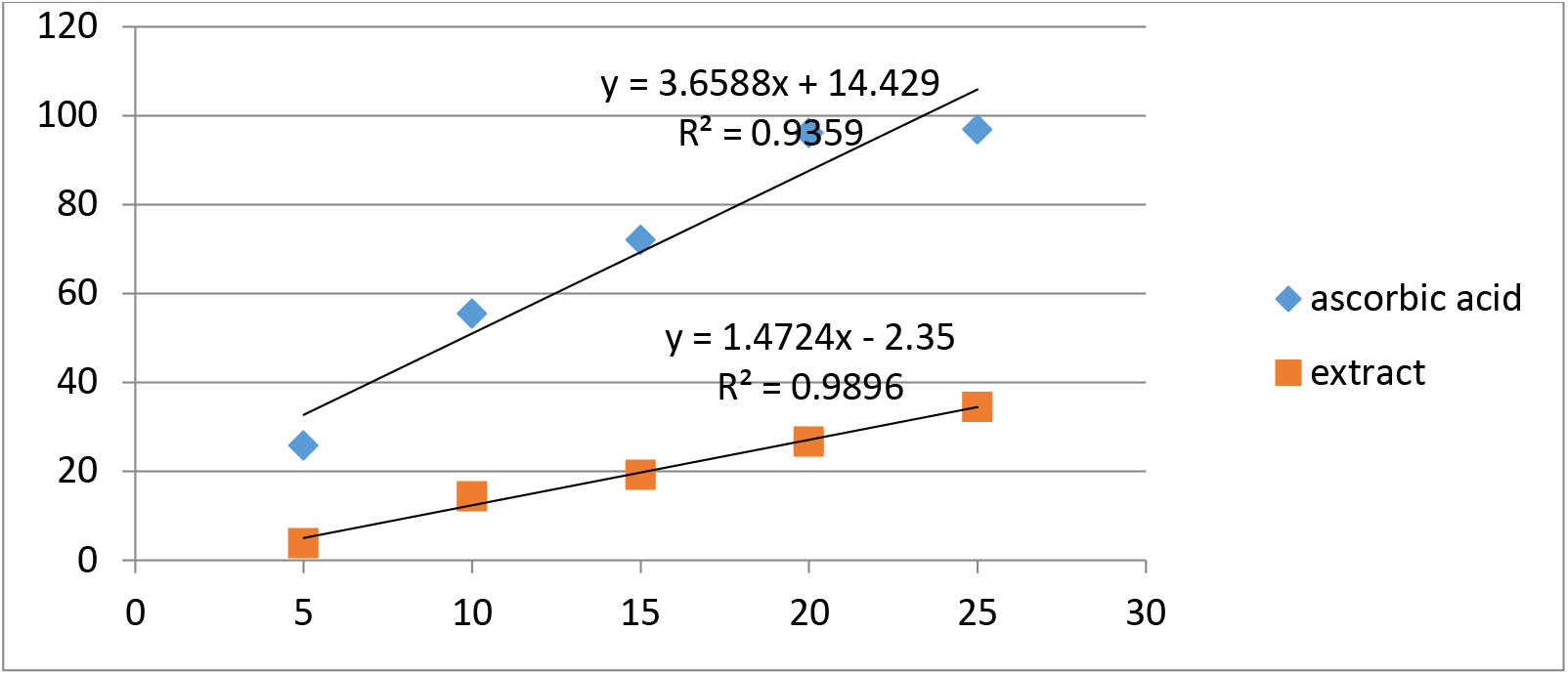
Regression line of ascorbic acid and extract

### Brine Shrimp Lethality Bioassay

The extract was also subjected to Brine Shrimp lethality bioassay for possible cytotoxic action. In this study the LC_50_ value of extract was found to be 312.5 μg/ml, whereas anticancer drug vincristine sulphate showed LC_50_ value of 7.33 μg/ml (Table 8). Whereas, previous study done by Ahmed et.al. showed the LC_50_ value of extract to be 400 μg/ml ^9^. According to Meyer, LC_50_ estimation of brine shrimp lethality assay, LC_50_ value of less than 1000 μg/ml is lethal while LC_50_ value more than 1000 μg/ml is deemed to be non-poisonous^26^.

**Table 8:** LC_50_ of the extract and standard Brine Shrimp Lethality Bioassay

| Test Sample | Concentration ($\mu\text{g/ml}$) | % mortality | $LC_{50}$ ($\mu\text{g/ml}$) |
| --- | --- | --- | --- |
| Vincristine sulphate | 0.25 | 20 | 7.33 |
|  | 0.5 | 30 |  |
|  | 1.0 | 30 |  |
|  | 5.0 | 40 |  |
|  | 10.0 | 60 |  |
| Extract | 50 | 20 | 312.5 |
|  | 100 | 30 |  |
|  | 200 | 40 |  |
|  | 400 | 70 |  |
|  | 800 | 90 |  |

### Synthesis of Zn-NPS

Reduction of zinc acetate to zinc oxide can be followed by color change. The formation of white precipitate indicated the formation of Zn-NPs.

### Characterization of NPs

#### 6.8.1 UV-Visible spectroscopy

The peak at 355nm indicates the formation of Zn-NPs.

**Figure 5:**
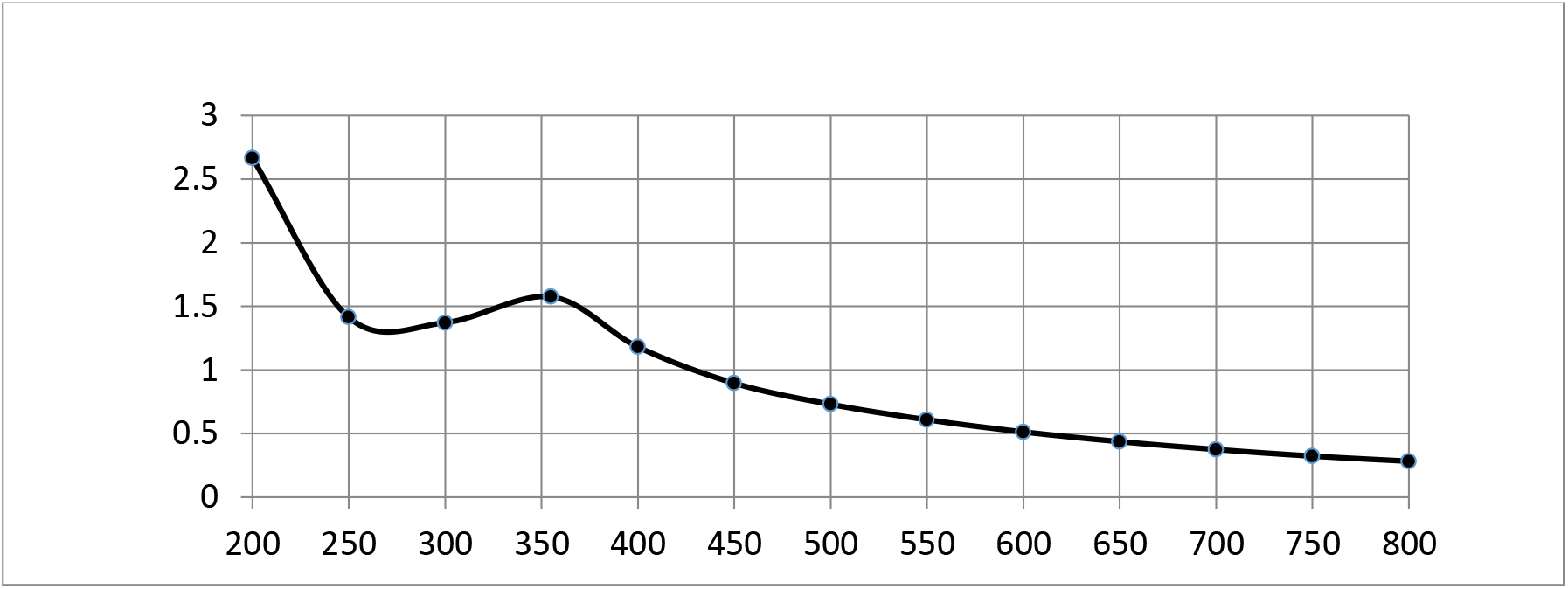
Peak showing maximum absorbance at 355nm

### X-ray diffraction

XRD peaks at 2Θ of 31.54, 34.216, 36.014, 47.32, 56.39, 62.70 and 67.80 can be attributed to the 100, 002, 101, 102, 110, 103 and 112 crystallographic planes of hexagonal wurtzite phase of ZnO, respectively. The XRD pattern thus clearly illustrated that Zn-NPs is formed in this study that are crystalline in nature. It also confirms the synthesized nanopowder was free of impurities as it does not contain any characteristics XRD peaks other than ZnO peaks^10^.

The size of crystal was calculated by Debye-Scherrer equation.

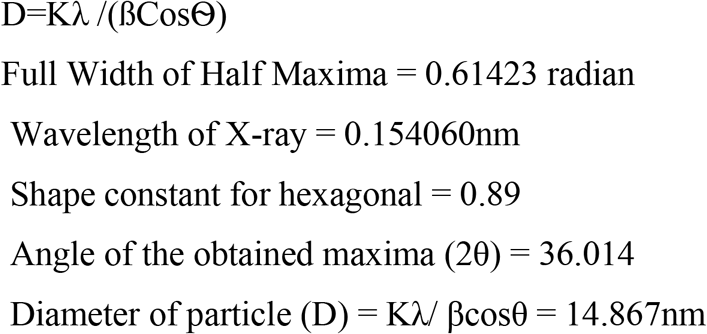

Therefore, the approximate average crystal size of zinc nanoparticle is 14.867nm

**Figure 6:**
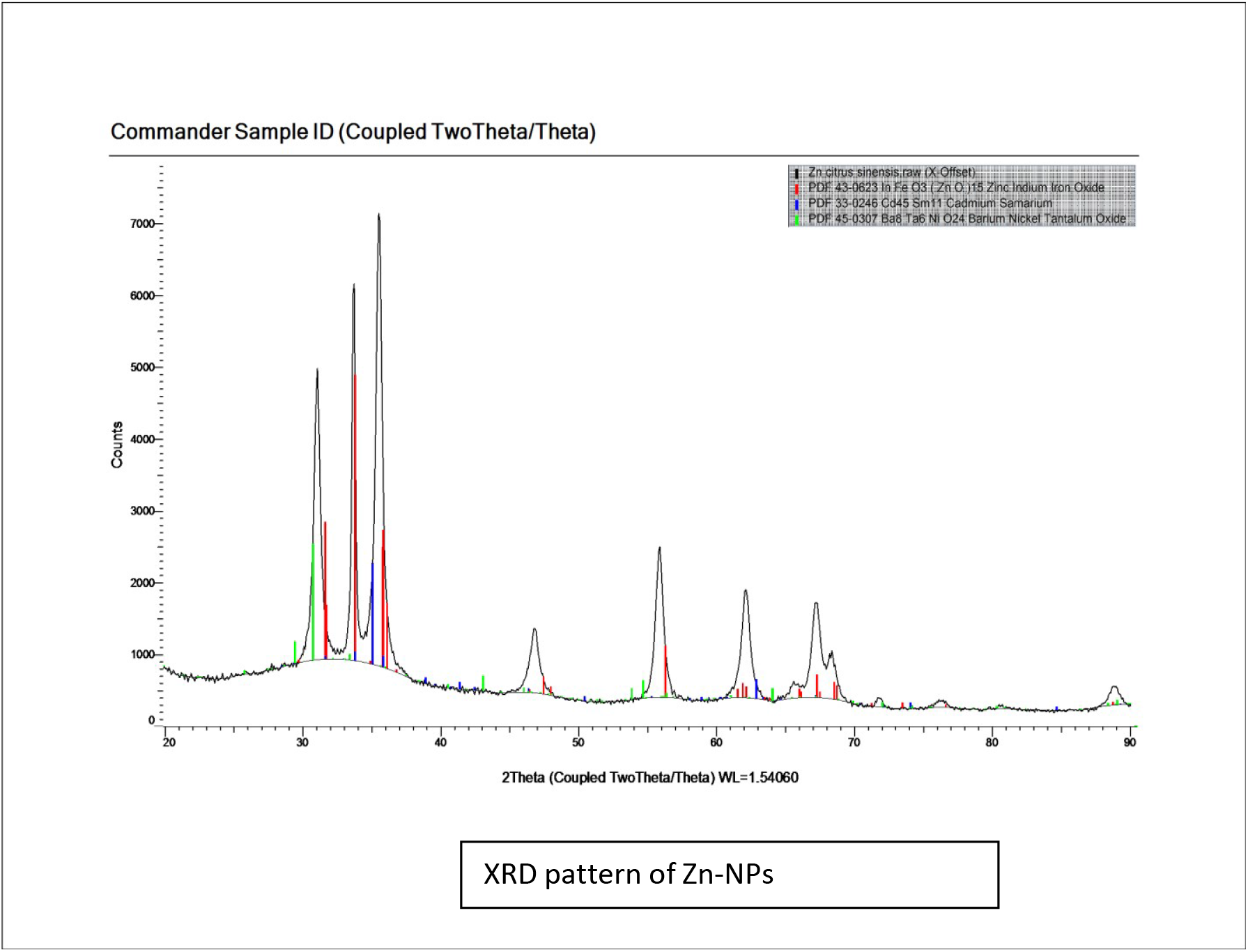
XRD pattern of ZN-NPs

#### 6.8.3 FTIR

The functional group present in extract responsible for reduction of zinc acetate to zinc oxide are:

**Table 9:**
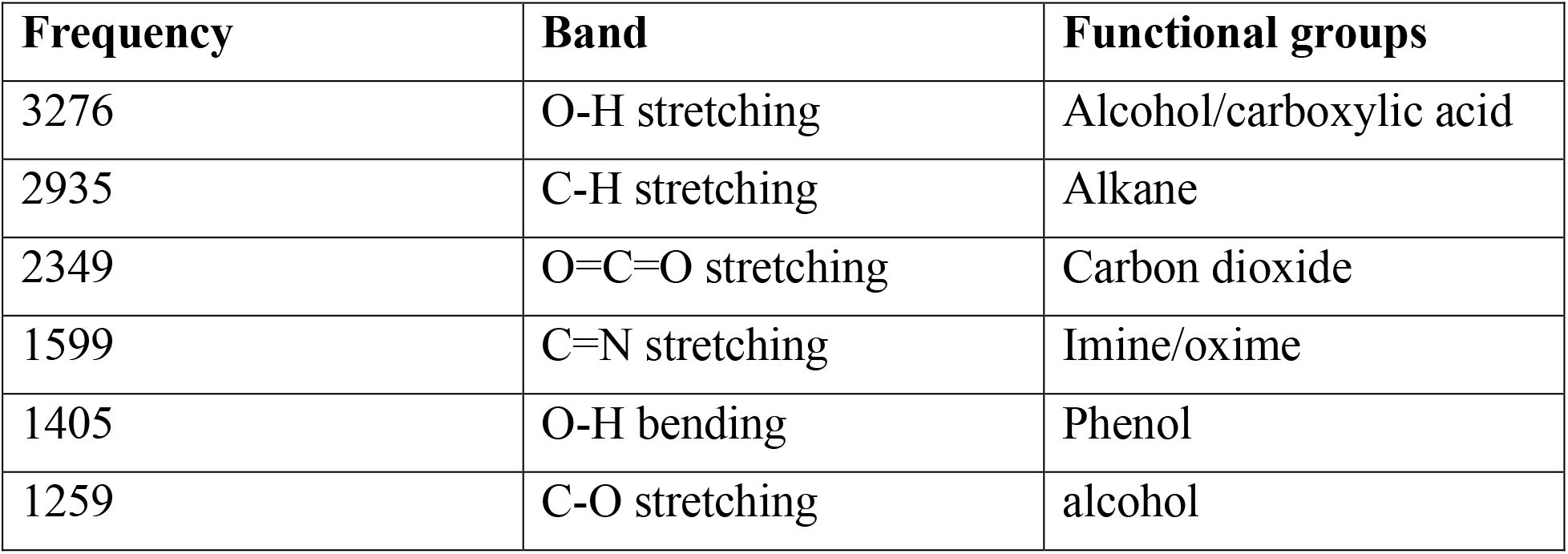
Functional group present in extract

**Figure 7:**
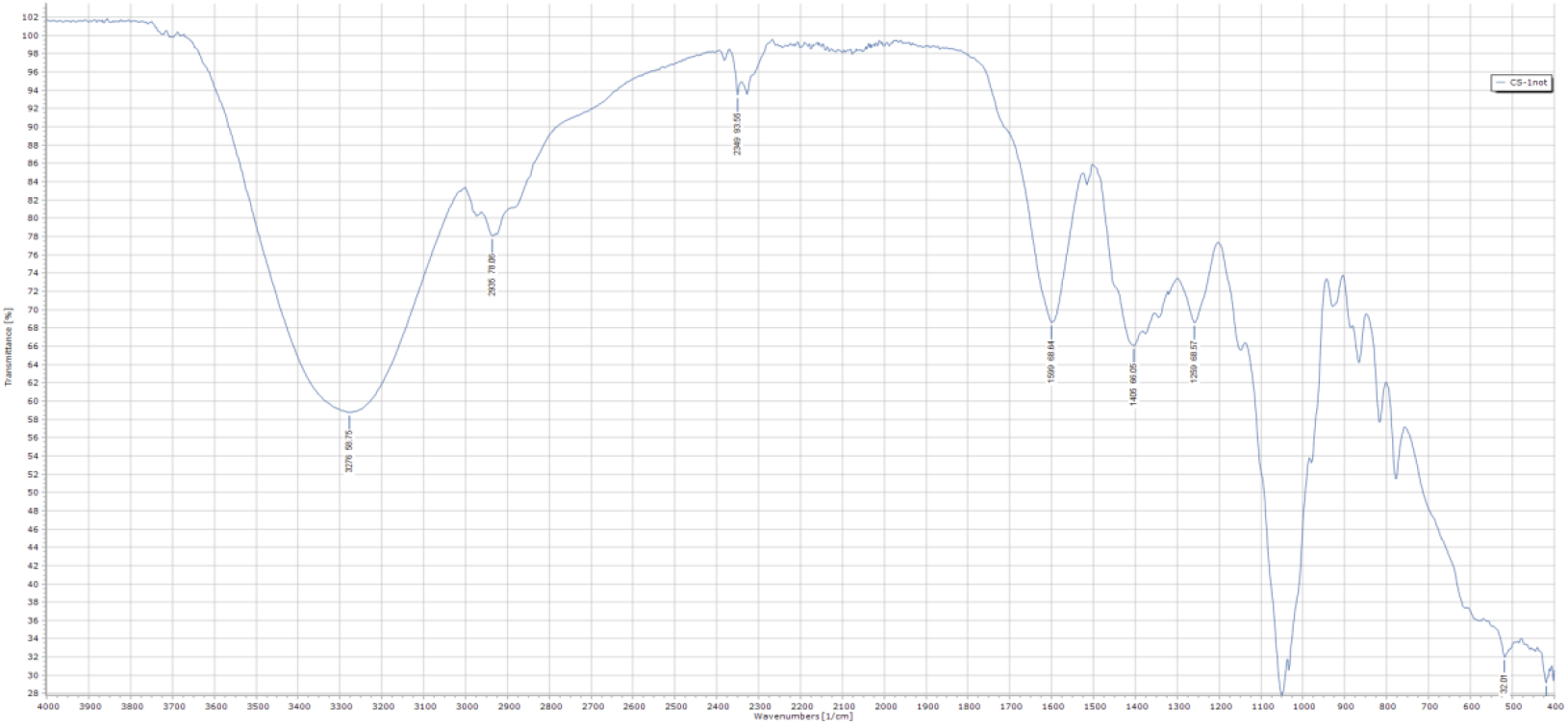
FTIR of extract The analysis was performed in a frequency range of 4000–400cm–1, at room temperature. The bands observed at 858cm-1 could be assigned to the C–H (aromatic) functional groups. The absorption peaks at 1377cm-1 can be attributed to the C–C stretching of aromatic rings. The spectrum presented a band at around 618cm-1; this signal is the characteristic bond signal of Zn–O, which confirms that the material is zinc oxide ^10^.

**Figure 8:**
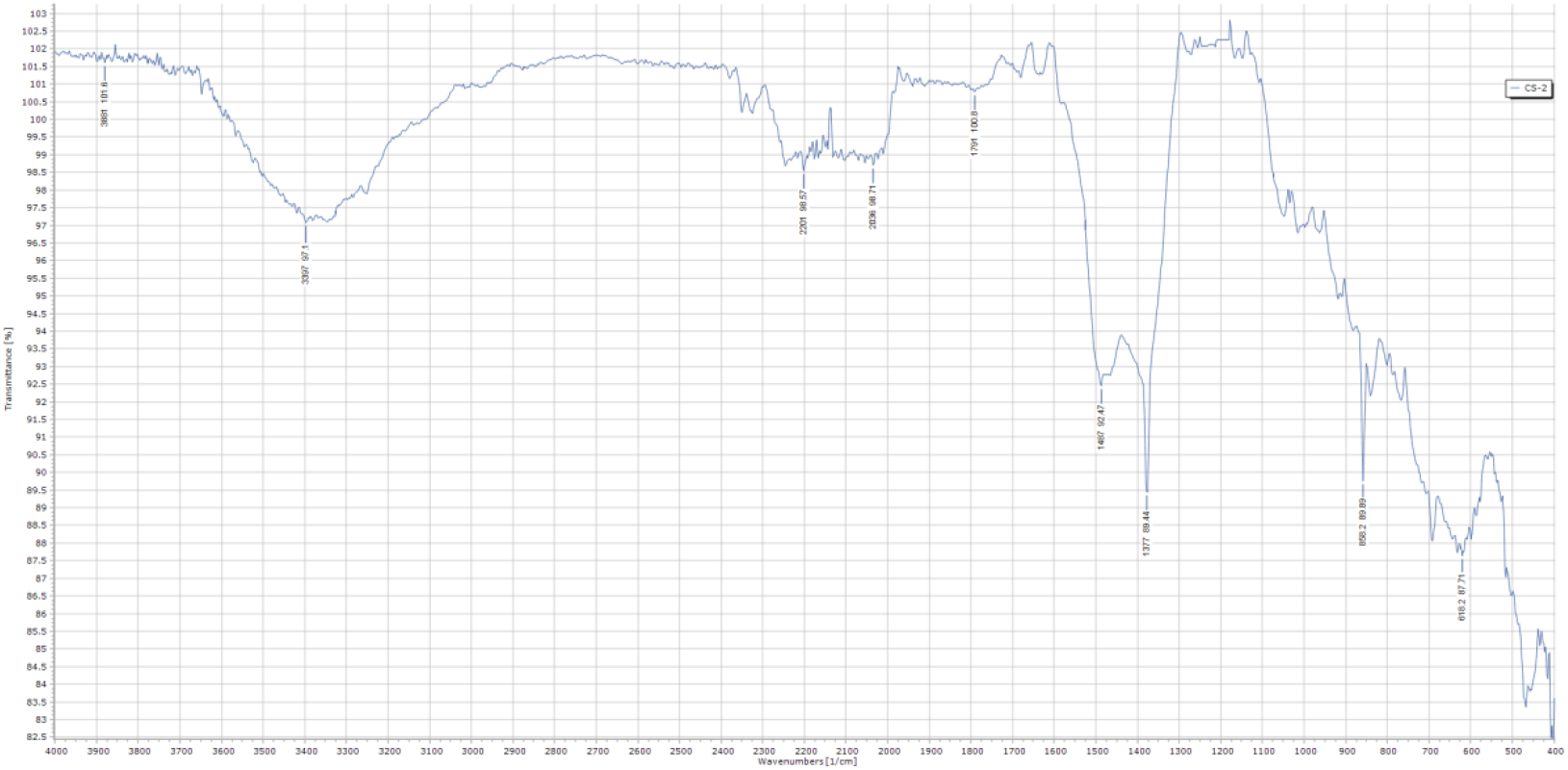
FTIR of Zn-NPs of *C. sinensis (L.) Osbeck*

### Antibacterial activity of the extract and nanoparticles

Table shows antibacterial activity of extract and Zn-NPs. The extract did not show any activity against *E. coli*.

**Table 10:** Antibacterial Activity of Extract and Zn-NPs

| Sample | Organisms | Zone of inhibition at diff. Concentration (mm) |  |  |  |
| --- | --- | --- | --- | --- | --- |
|  |  | 40µg/ml | 80µg/ml | 160µg/ml | 320µg/ml |
| Extract | *K. pneumoniae* | 2 | 4 | 4 | 4 |
|  | *E. coli* | 0 | 0 | 0 | 0 |
|  | *B. subtilis* | 2 | 4 | 5 | 5 |
|  | *S. aureus* | 0 | 0 | 4 | 5 |
| Zn-NPs | *K. pneumoniae* | 3 | 3 | 5 | 6 |
|  | *E. coli* | 3 | 3 | 5 | 7 |
|  | *B. subtilis* | 5 | 7 | 7 | 9 |
|  | *S. aureus* | 2 | 3 | 5 | 7 |

**Table 11:** Antibacterial activity of Standard

| Standard | Organism | Zone of inhibition at diff. Concentration (mm) |  |  |  |
| --- | --- | --- | --- | --- | --- |
|  |  | 40µg/ml | 80µg/ml | 160µg/ml | 320µg/ml |
| Gentamycin | *K. pneumoniae* | 3 | 3 | 4 | 7 |
|  | *E. coli* | 8 | 9 | 13 | 15 |
| Azithromycin | *B. subtilis* | 10 | 12 | 19 | 21 |
|  | *S. aureus* | 5 | 7 | 12 | 14 |

**Table 12:** Antibacterial activity of control

| Organisms | Zone of inhibition (mm) |  |  |
| --- | --- | --- | --- |
|  | Zinc acetate<br>(0.02) | Distilled water | DMSO |
| *K. pneumoniae* | 0 | 0 | 0 |
| *E. coli* | 0 | 0 | 0 |
| *B. subtilis* | 0 | 0 | 0 |
| *S. aureus* | 3 | 0 | 0 |

## Acknowledgements

We would like to thank Mrs. Deepti Piya Baniya, Mrs. Mijala Bajracharya, Mrs. Sarita Khanal, Mr. Bechan Raut, and Mr. Bishwo Raj Dhungana at Manmohan Institute of Health Sciences, Kathmandu.

## References

1. Khan, I., Saeed, K., & Khan, I. (2017, May 18). Nanoparticles: Properties, applications and toxicities. https://doi.org/10.1016/j.arabjc.2017.05.011

2. Singh, O. V. (2015). Bio-nanoparticles: Biosynthesis and sustainable biotechnological implications. Hoboken, NJ: Wiley-Blackwell.

3. Mosquera, J., García, I., & Liz-Marzán, L. M. (2018). Cellular Uptake of Nanoparticles versus Small Molecules: A Matter of Size. Accounts of Chemical Research, 51(9), 2305–2313. doi:DOI: 10.1021/acs.accounts.8b00292

4. Ahmed, S., & Ikram, S. (2015). Synthesis of Gold Nanoparticles using Plant Extract: An Overview. Nano Research & Applications, 1(1).

5. Tripathi, D. K., Ahmad, P., Kumar, D., Nawal, C., Dubey, K., & Sharma, S. (2017). Nanomaterials in Plants, Algae, and Microorganisms (First ed., Vol. 1). London: Academic Press.

6. Zhang, X., Liu, Z., Shen, W., & Gurunathan, S. (2016). Silver Nanoparticles: Synthesis, Characterization, Properties, Applications, and Therapeutic Approaches. International Journal of Molecular Sciences, 17(1354). doi:10.3390/ijms17091534

7. Punjabi, K., Mehta, S., Chavan, R., Chitalia, V., Deogharkar, D., & Deshpande, S. (2018). Efficiency of Biosynthesized Silver and Zinc Nanoparticles Against Multi-Drug Resistant Pathogens. Frontiers in Microbiology, 9(2207). doi:https://doi.org/10.3389/fmicb.2018.02207

8. Musa, D. D., Sangodele, F., & Hafiz, S. S. (2019). PHYTOCHEMICAL ANALYSIS AND ANTIBACTERIAL ACTIVITY OF ORANGE (Citrus sinensis) PEEL. FUDMA Journal of Sciences (FJS), 3(1), 375–380.

9. Ahmed, N., Ahmed, T., Akbar, N., & Ahmed, S. (2016). PHYTOCHEMICAL SCREENING, CYTOTOXIC AND HYPOGLYCEMIC ACTIVITY OF METHANOLIC EXTRACT OF CITRUS SINENSIS PEEL. NTERNATIONAL RESEARCH JOURNAL OF PHARMACY, 7(3), 43–48. doi:DOI: 10.7897/2230-8407.07328

10. Nava, O., Soto-Roblesa, C., Gomez-Gutierrez, C., Vilchis-Nestor, A., Castro-Beltran, A., Olivas, A., & Luque, P. (2017). Fruit peel extract mediated green synthesis of zinc oxide nanoparticles. Journal of Molecular Structure, 1147, 1–5. http://dx.doi.org/10.1016/j.molstruc.2017.06.078

11. Dobrucka, R., & Długaszewska, J. (2016). Biosynthesis and antibacterial activity of ZnO nanoparticles using Trifolium pratense flower extract. Saudi Journal of Biological Sciences Pages, 23(4), 517–523. doi:https://doi.org/10.1016/j.sjbs.2015.05.016

12. Kokate, C. K. (2014). Practical Pharmacognosy (Fifth ed.). Delhi: Vallabh Prakashan.

13. Harborne, A. J. (1998). Phytochemical Methods A Guide to Modern Techniques of Plant Analysis (Second ed.). London: Chapman and Hall.

14. Singleton, V. L., Orthofer, R., & Lamuela-Raventós, R. M. (1999). Analysis of total phenols and other oxidation substrates and antioxidants by means of folin-ciocalteu reagent. Methods in Enzymology, 299, 152–178. doi:https://doi.org/10.1016/S0076-6879(99)99017-1

15. Silva, L. A., Pezzini, B. R., & Soares, L. (2015). Spectrophotometric determination of the total flavonoid content in Ocimum basilicum L. (Lamiaceae) leaves. Pharmacogn Mag., 11(41), 96–101. doi:doi: 10.4103/0973-1296.149721

16. Liang, N., & Kitts, D. D. (2019). Antioxidant property of coffee components: Assessment of methods that define mechanisms of action. Multidisciplinary Digital Publishing Institute, 19(11). doi:DOI: 10.3390/molecules191119180

17. Mahdi-Pour, B., Jothy, S. L., Latha, L. Y., Chen, Y., & Sasidharan, S. (2012). Antioxidant activity of methanol extracts of different parts of Lantana camara. Asian Pac J Trop Biomed., 2(12), 960–965. doi:doi: 10.1016/S2221-1691(13)60007-6

18. Finney, D. J. (1971). Probit Analysis (3rd ed.). New York: Cambridge University Press. doi:https://doi.org/10.1002/jps.2600600940

19. Janjal, S., Rajbhoj, A., & Gaikwa, S. (2017). Synthesis and characterization of zinc oxide nanoparticles using green method. Research Journal of Chemical Sciences, 7(4), 40–42.

20. Osman, D. A., & Mustafa, M. A. (2015). Synthesis and Characterization of Zinc Oxide Nanoparticles using Zinc Acetate Dihydrate and Sodium Hydroxide. Journal of Nanoscience and Nanoengineering, 1(4), 248–251.

21. Selim, Y. A., Azb, M. A., Ragab, I., & El-Azim, M. H. (2020). Green Synthesis of Zinc Oxide Nanoparticles Using Aqueous Extract of Deverra tortuosa and their Cytotoxic Activities. Sci Rep, 10. doi:https://doi.org/10.1038/s41598-020-60541-1

22. He, K., Chen, N., Wang, C., Wei, L., & Chen, J. (2018). Method for Determining Crystal Grain Size by X-Ray Diffraction. Crystal Research and Technology, 53(2). doi:DOI: 10.1002/crat.201700157

23. Bonev, B., Hooper, J., & Parisot, J. (2008). Principles of assessing bacterial susceptibility to antibiotics using the agar diffusion method. Journal of Antimicrobial Chemotherapy, 6(61), 1295–1301. doi:DOI: 10.1093/jac/dkn090

24. Liew, S. S., Ho, W. Y., Yeap, S. K., & Sharifudin, S. A. (2018). Phytochemical composition and in vitro antioxidant activities of Citrus sinensis peel extracts. PeerJ, 6. doi:https://doi.org/10.7717/peerj.5331

25. Oladele, O. O., & Aborisade, A. T. (2018). Activity and characterization of antifungal compounds from the peel of sweet orange Citrus sinenesis fruits. J. Crop Prot., 7(3), 327–335.

26. Meyer, B. N., Ferrigni, N. R., Putnam, J. E., Jacobsen, L. B., Nichols, D. E., & McLaughlin, J. L. (1982). Brine shrimp: A convenient general bioassay for active plant constituents. Planta Medica, 45(5), 31–34. doi:DOI: 10.1055/s-2007-971236

